# GenEpi: Gene-based Epistasis Discovery Using Machine Learning

**DOI:** 10.1101/421719

**Authors:** Yu-Chuan Chang, June-Tai Wu, Ming-Yi Hong, Yi-An Tung, Ping-Han Hsieh, Sook Wah Yee, Kathleen M. Giacomini, Yen-Jen Oyang, Chien-Yu Chen, for the Alzheimer’s Disease Neuroimaging Initiative

## Abstract

Genome-wide association studies (GWAS) provide a powerful means to identify associations between genetic variants and phenotypes. However, GWAS techniques for detecting epistasis, the interactions between genetic variants associated with phenotypes, are still limited. We believe that developing an efficient and effective GWAS method to detect epistasis will be a key for discovering sophisticated pathogenesis, which is especially important for complex diseases such as Alzheimer’s disease (AD). In this regard, this study presents GenEpi, a computational package to uncover epistasis associated with phenotypes by the proposed machine learning approach. GenEpi identifies both within-gene and cross-gene epistasis through a two-stage modeling workflow. In both stages, GenEpi adopts two-element combinatorial encoding when producing features and constructs the prediction models by L1-regularized regression with stability selection. The simulated data showed that GenEpi outperforms other widely-used methods on detecting ground-truth epistasis. As real data is concerned, this study uses AD as an example to reveal the capability of GenEpi in finding disease-related variants and variant interactions that show both biological meanings and predictive power. **Availability:** GenEpi is an open-source python package and available free of charge only for non-commercial users. The package can be downloaded from https://github.com/Chester75321/GenEpi, and has also been published on The Python Package Index.

## 1. Introduction

Genome-wide association studies (GWAS) is a univariate examination of a genome-wide set of genetic variants to determine if any single variant is associated with the phenotype of interest [1]. The first GWAS was published in 2002 [2], and three years later, the most remarkable GWAS regarding age-related macular degeneration (AMD) was published [3]. Their study investigated the association of 105,980 single nucleotide polymorphisms (SNPs) with AMD on 96 cases and 50 control subjects. This study showed that the SNPs in the complement factor H (CFH) gene, including a non-synonymous SNP, are significantly associated with AMD. Up to 2019, there have been more than hundreds of thousands individuals being studied in typical GWAS protocols, and over 210,498 variant-disease associations between 117,337 SNPs and 10,358 phenotypes have been discovered [4]. These studies demonstrated the potential of GWAS to identify genetic variants associated with many categories of phenotypes, including risk for diseases such as various cancers, and variation in therapeutic and adverse response to drugs. However, the success of univariate GWAS is limited to monogenic phenotypes (e.g. Mendelian diseases). The impact of variant interactions, also known as epistasis on the formation of diseases [5] is often underestimated in traditional GWAS analysis [6–8].

A major limitation of traditional GWAS is that it considers only one genetic variant at a time, and ignores underlying epistasis of variants that might have stronger associations [9]. Researchers have found that GWAS has limitation in identifying the association in complex diseases [10, 11]. Easton et al. suggested that a number of susceptible loci identified by GWAS usually have very small effect sizes [12]. Studies have also demonstrated that the existence of epistasis is an important factor contributing to phenotypes, especially in complex diseases such as hypertension, diabetes and obesity [11]. Therefore, developing analytical methods to identify epistasis efficiently is critical to understanding the genetic factors [8, 13], and has attracted a wide range of research interests in recent years [7, 14].

There are, however, two main challenges to discover epistasis: computational complexity and statistical power [15]. The first challenge results from the curse of dimensionality. When more genetic variants are considered, the number of interactions increases exponentially. Based on the specification of a major commercial technology, Illumina Arrays, a whole-genome array can investigate over 4 million markers per sample simultaneously. In order to evaluate the pairwise interactions from this microarray, about 8×10^12^ statistical tests need to be processed. Even though Marchini et al. have demonstrated that pairwise interactions of 3×10^5^ loci is computationally possible with currently available computational resources, it still remains challenging when the Illumina Arrays are considered [16]. The second challenge is the issue of statistical power. Since a huge number of statistical tests are conducted on a limited sample size with high-dimensional interactions, many false positives arise by random chances. In these regards, new methods have been developed [11, 17]. Statistical approaches include FastEpistasis [18] and BOOST [19]; both of them has been included in a well-known GWAS software called PLINK [20, 21]. Machine learning approaches such as Multifactor Dimensionality Reduction [22], ReliefF [23], random forest-like algorithms [24–26] and other methodologies have also been developed for detecting epistasis [17].

Since the biological experiments used to validate these methodologies such as the AMD in traditional GWAS are still in demand, there are no standard analysis methods for epistasis despite the rapid improvement in computational performance.In 2016, Murk used FastEpistasis and BOOST to search SNP-SNP interactions on a huge dataset called Genetic Epidemiology Research on Adult Health and Aging (GERA) that included 78,486 subjects, but still failed to detect a significant and replicable interaction after exhaustively searching through 45 billion possible interactions for 10 complex diseases of interest [27]. Alzheimer’s disease (AD) is one of the most important complex diseases and its pathogenesis, which clearly has a genetic basis, is still ill-defined. In 2014, Sage Bionetworks held a competition called The Dialogue for Reverse Engineering Assessments and Methods Challenge (DREAM Challenge) for AD, which tried to use crowdsourcing to assess the capability of current computational methods to predict the change in cognitive examination based on genetic data. However, no significant contribution of genetic features except the *APOE* haplotype to the predictive performance was observed by any competition teams [28]. To address this problem, this study presents GenEpi, a package to reveal epistasis related to the phenotype using machine learning and introduces the application of GenEpi on predicting the diagnosis of AD.

## 2. Results

This study compared GenEpi with several commonly used algorithms for detecting epistasis, including FastEpistasis, BOOST and ReliefF. The simulation data demonstrated that GenEpi outperforms the other methods in ranking the true epistasis as the top one. As real data is concerned, the results suggest that the epistasis selected by GenEpi has the best predictive power for diagnosis of AD. The proposed model of predicting AD contains 14 genetic features, including 24 SNPs from 12 genes that contain the well-known causal gene, APOE. The 2-fold cross validation (CV) and leave-one-out CV (LOO CV) accuracy of this model are 0.829 and 0.832, respectively. The results on AD demonstrated that GenEpi has the ability to detect the epistasis associated with the phenotype effectively and efficiently. The released package can be generalized to largely facilitate the studies of many complex diseases in the near future.

We will demonstrate our experiments in following three parts of this section. In the first part, we applied GenEpi and other algorithms for detecting epistasis, including FastEpistasis [18], BOOST [19] and ReliefF [23, 29] on simulation data for validation and comparison. In the second part, we applied GenEpi on the ADNI dataset to categorize each sample as control subjects or AD patients, evaluated by precision, recall, accuracy and F1 score (2 × (precision × recall) / (precision + recall)). In the final part, we compared GenEpi with other algorithms on the ADNI dataset in terms of computing time and prediction performance on real data.

### Experiments on simulation data

We applied different algorithms on simulation data for validation and comparison. All of the simulation datasets are generated by the simulator GAMETES [30], which is publicly accessible on the web site https://popmodels.cancercontrol.cancer.gov/gsr/packages/gametes/. We designed two types of simulation datasets: basic and complex models. The ‘Model 1’, ‘Model 2’ and ‘Model 3’ are simulation datasets with the basic model, which means that each dataset contained only one epistasis consisting of a SNP pair. All of the basic-model datasets are in the same setting as follows: #individuals = 2000, case/control ratio = 1, #SNPs = 100, #replicates = 100, minor allele frequency of target SNPs = 0.2, and heritability = 0.2. The complex model means one dataset contains multiple epistasis from different SNP pairs. Here, the ‘Combined Model 1+2+3’ is a complex model dataset containing three epistasis from the previous three basic models.

Fig. 1 provided the results of these four simulation datasets. Fig. 1(a) shows that the ranking of the target epistasis reported by GenEpi in the 100 replicates of each basic-model dataset are always ranked as the first. In contrast, for FastEpistasis and BOOST, the medians of the ranking of the target epistasis among the 100 runs of simulation are one but the averages are not. The number of failures of FastEpistasis and BOOST in 100 replicates of three basic models are 6, 1, 16 and 5, 1, 14, respectively. For the result of the complex model dataset in Fig. 1(b), the superiority of GenEpi over other algorithms is more obvious. In the 100 runs of simulation, GenEpi reported the three target epistasis as the top three important features every time. In contrast, FastEpistasis and Boost failed to report the three target epistasis as the top three important features consistently.

**Fig. 1.**
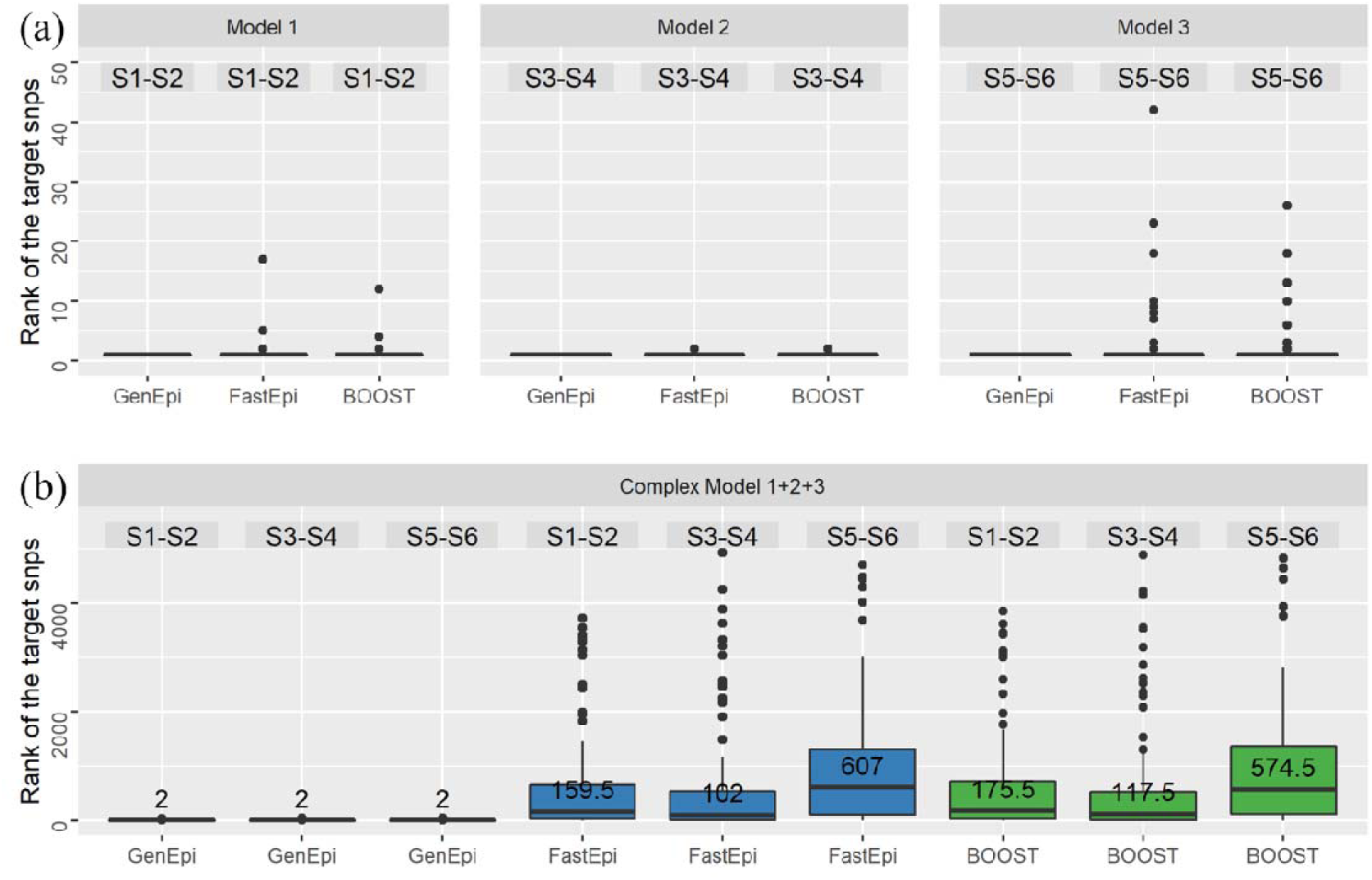
The boxplot for the rank of the target epistasis in different algorithms. (a) The results of three basic-model datasets with one epistasis consisting of a SNP pair. (b) The result of the complex-model dataset, which contained three epistasis. The ‘S1-S2’ means the epistasis between SNP 1 and SNP 2 and so on. The values on the boxplot are the medians of the rank of the target epistasis among the 100 runs of simulation.

When ReliefF was compared, since the Python package scikit-rebate [29] that we used for implementing ReliefF only reports the importance of individual SNPs instead of the scores for epistasis (SNP pairs), we listed the medians of ReliefF’s ranking for each SNP in the target epistasis in Table 1. Table 1 reveals that ReliefF can detect the SNPs in the target epistasis in the basic models, but failed to report the three target epistasis as the top three important features in the complex model dataset.

**Table 1.** The medians of the rank of the SNPs in the target epistasis for ReliefF

|  | SNP 1 | SNP 2 | SNP 3 | SNP 4 | SNP 5 | SNP 6 |
| --- | --- | --- | --- | --- | --- | --- |
| Basic Model | 1 | 2 | 1 | 2 | 1 | 2 |
| Complex Model | 7 | 8.5 | 9.5 | 11.5 | 11 | 19.5 |

The superiority of GenEpi is owing to the proposed two-element combinatorial encoding of the genotype features and the L1-regularized regression with stability selection. In contrast with other statistical algorithms such as FastEpisasis and BOOST, which only evaluate the epistasis of a SNP pair one at a time, GenEpi considers interactions between combinatorial features by multivariate models. Moreover, the false positives among the epistasis can be filtered out by resampling and remodeling the dataset hundreds of times. To evaluate the effect of stability selection, we applied both L1-regularized regression with and without stability selection on the complex model dataset to compare the number of false positives, which is defined as the number of non-target epistasis in the final output of GenEpi. As shown in Fig. 2, stability selection can reduce the mean false positive rate effectively and minimize the variance of false positive rate as well.

**Fig. 2.**
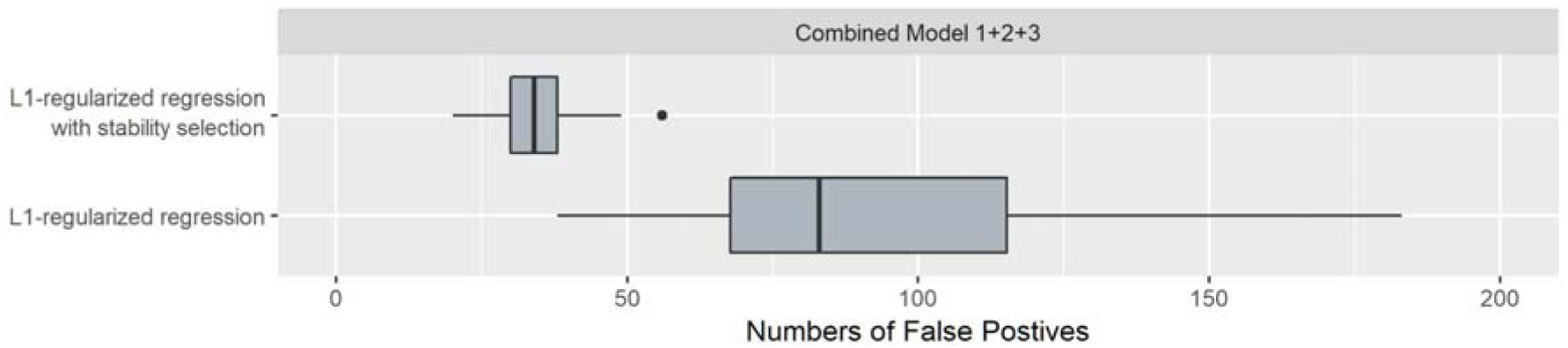
The boxplot of false positives in L1-regularized regression with and without stability selection.

### Classifying AD patients

In predicting control subjects or AD patients, we applied GenEpi on the 364 samples with CN (as control) or AD. After dimensionality reduction, 12,102,888 out of the 12,809,667 SNPs in the ADNI dataset were retained, and 4,916,249 of them are located in 20,206 genes (Table 2). In the step 4 of selecting epistasis, there are 34,689 genetic features selected and 765 of them are single SNP features, while the other 33,924 are epistasis features within genes. The final model contained 14 genetic features, including 24 SNPs from 12 genes. These features contained two single SNP features, 11 within-gene epistasis features and one cross-gene epistasis feature. As shown in Table 3, the 2-fold cross validation (CV) and leave-one-out CV (LOO CV) accuracy of this model are 0.829 and 0.832, respectively.

**Table 2.**
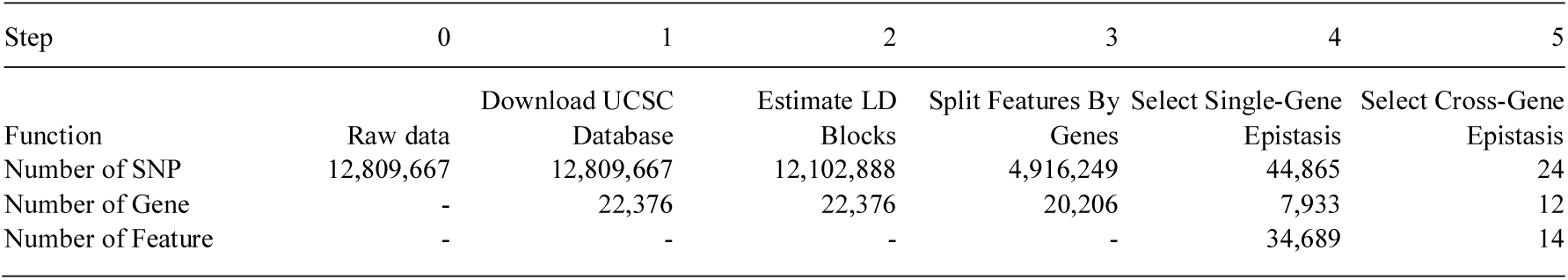
The summary of the number of SNPs, genes and features in each step of GenEpi.

**Table 3.** The score of different models in predicting control subjects or AD patients.

|  | Precision | Recall | F1 Score | Accuracy |
| --- | --- | --- | --- | --- |
| Training | 0.9633 | 0.8537 | 0.9052 | 0.9396 |
| 2-fold CV | 0.7748 | 0.6992 | 0.7350 | 0.8297 |
| LOO CV | 0.8100 | 0.6585 | 0.7265 | 0.8324 |

F1 Score = $2 \times (\text{Precision} \times \text{Recall}) / (\text{Precision} + \text{Recall})$ ; ‘Training’ stands for the process of a single-loop CV; ‘2-fold CV’ means that
2-fold CV was used in the external loop of double CV; ‘LOO CV’ means that LOO CV was used in the external loop of double CV.

We listed the statistical significance of the selected genetic features in Table 4. The first column lists each feature by its RSID (Reference SNP cluster ID) and the genotype (denoted as RSID_genotype), the pairwise epistasis features are represented using two SNPs. The last column describes the genes where the SNPs are located according to the genomic coordinates. We used a star sign to denote the epistasis between genes. Here, only the feature (rs3130614_BB, rs41276317_AB) is cross-gene epistasis (for MICB and TOB2). The weights in the second column were extracted from the linear model we defined in Section 2.4. The signs of the weights indicate if a feature is a causal or protective genotype, which is consistent with the corresponding odds ratio. The p-value of the χ^2^ test showed that these features are significantly associated with the phenotype.

**Table 4.**
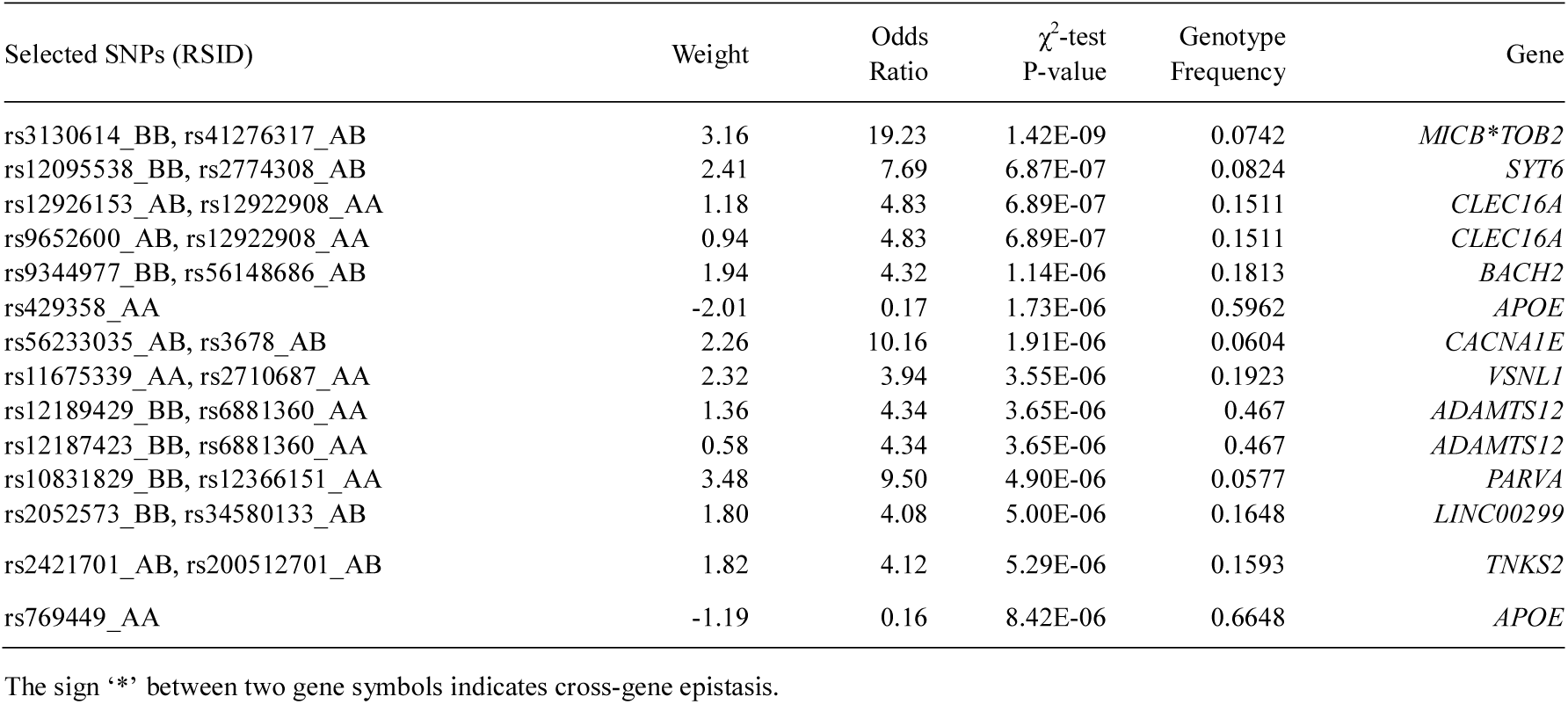
The statistical significance of genetic features selected by GenEpi in predicting patients with AD.

| Selected SNPs (RSID) | Weight | Odds Ratio | $\chi^2$-test<br>P-value | Genotype<br>Frequency | Gene |
| --- | --- | --- | --- | --- | --- |
| rs3130614_BB, rs41276317_AB | 3.16 | 19.23 | 1.42E-09 | 0.0742 | *MICB\*TOB2* |
| rs12095538_BB, rs2774308_AB | 2.41 | 7.69 | 6.87E-07 | 0.0824 | *SYT6* |
| rs12926153_AB, rs12922908_AA | 1.18 | 4.83 | 6.89E-07 | 0.1511 | *CLEC16A* |
| rs9652600_AB, rs12922908_AA | 0.94 | 4.83 | 6.89E-07 | 0.1511 | *CLEC16A* |
| rs9344977_BB, rs56148686_AB | 1.94 | 4.32 | 1.14E-06 | 0.1813 | *BACH2* |
| rs429358_AA | -2.01 | 0.17 | 1.73E-06 | 0.5962 | *APOE* |
| rs56233035_AB, rs3678_AB | 2.26 | 10.16 | 1.91E-06 | 0.0604 | *CACNA1E* |
| rs11675339_AA, rs2710687_AA | 2.32 | 3.94 | 3.55E-06 | 0.1923 | *VSNL1* |
| rs12189429_BB, rs6881360_AA | 1.36 | 4.34 | 3.65E-06 | 0.467 | *ADAMTS12* |
| rs12187423_BB, rs6881360_AA | 0.58 | 4.34 | 3.65E-06 | 0.467 | *ADAMTS12* |
| rs10831829_BB, rs12366151_AA | 3.48 | 9.50 | 4.90E-06 | 0.0577 | *PARVA* |
| rs2052573_BB, rs34580133_AB | 1.80 | 4.08 | 5.00E-06 | 0.1648 | *LINC00299* |
| rs2421701_AB, rs200512701_AB | 1.82 | 4.12 | 5.29E-06 | 0.1593 | *TNKS2* |
| rs769449_AA | -1.19 | 0.16 | 8.42E-06 | 0.6648 | *APOE* |

The sign ‘\*’ between two gene symbols indicates cross-gene epistasis.

### Comparison with different algorithms

In this section, we compared GenEpi with other algorithms for detecting epistasis, including FastEpistasis [18], BOOST [19] and ReliefF [23] in terms of computing time and prediction performance. We used Microsoft Azure E32 v3 as the computing resource, which contains 32 CPUs and 256 GB RAM. Since the PLINK (version 2.0) has imported FastEpistasis and BOOST, we used PLINK to test these two algorithms. For ReliefF, we employed a Python package called scikit-rebate [29] for implementation. Among these algorithms, only FastEpistasis can afford the computation of the whole set of SNPs. In this regard, 12,809,667 SNPs were used by FastEpistasis (Table 5). On the other hand, GenEpi only focuses on the SNPs in the gene regions. In this regard, the number of input SNPs for estimating epistasis reduced to 4,916,249. BOOST took the same subsets of SNPs as GenEpi (Table 5). When taking the same subset of SNPs as GenEpi and BOOST, ReliefF still caused memory errors.

**Table 5.** The comparison of different algorithms.

| Algorithm | # Input SNP | Time Cost | Top 15 | Top 30 | Top 45 | Top 60 |
| --- | --- | --- | --- | --- | --- | --- |
| GenEpi | 4,916,249 | 9.95 | 0.76 | 0.72 | 0.71 | 0.68 |
| BOOST | 4,916,249 | 2157.6 | 0.31 | 0.24 | 0.30 | 0.37 |
| ReliefF | 33,868 | 0.11 | 0.52 | 0.48 | 0.45 | 0.46 |
| FastEpistasis | 12,809,667 | 836.8 | 0.62 | 0.61 | 0.60 | 0.59 |

‘Time Cost’ is the time spent on identifying the epistasis, which was measured by single CPU time in days. The values in column top 15, top 30, top 45 and top 60 are the 2-fold CV scores. The 2-fold CV scores are the F1 scores.

Therefore, we used the subsets of SNPs that selected by STAGE 1 of GenEpi as the input of ReliefF, which are 33,868. We selected the top 15, 30, 45 and 60 rankings from the results of these algorithms for comparing the prediction performance, and used L1-regularized regression to build the models for classifying AD patients for comparison. Table 5 show that GenEpi is the most efficient method, which only cost 9.95 CPU-days. Moreover, GenEpi had the best prediction performance despite the fact that GenEpi only uses the subset of SNPs from the final model. GenEpi shows that the time needed for identifying epistasis can be drastically reduced, without compromise to the performance. We provided the ROC curves for the classification task in Fig. 3, and it shows that GenEpi achieved the best performance in double 2-fold CV procedures, of which the area under the curve (AUC) is 0.85.

**Fig. 3.**
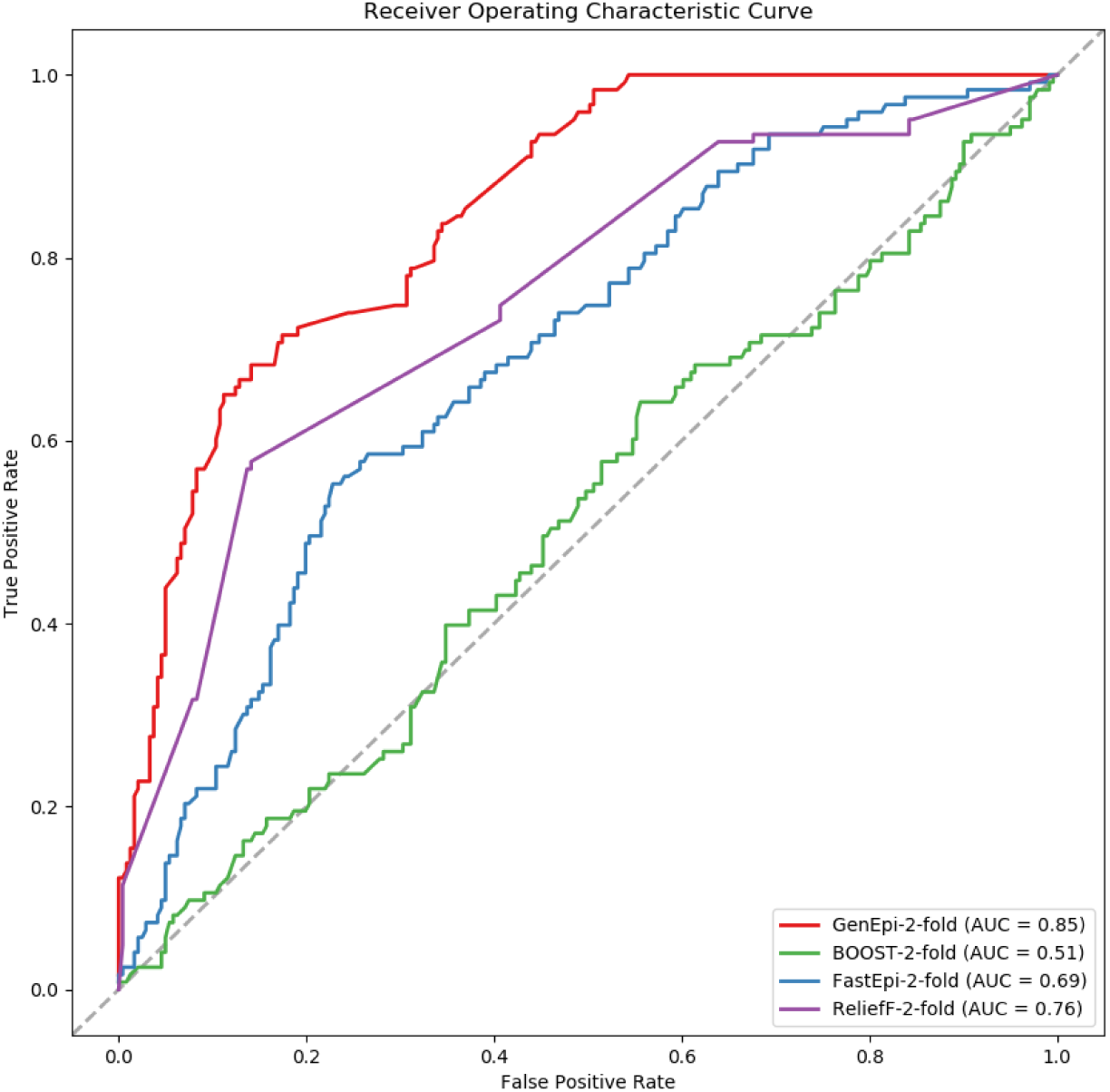
The ROC curves of different algorithms.

## 3. Discussion

The results in the previous section revealed the power of GenEpi to identify phenotype-associated epistasis efficiently. GenEpi selected 14 features from 12 genes to categorize patients with AD. Since AD is a chronic neurodegenerative disease, our findings would be supported if the gene identified by GenEpi are expressed in brains. We downloaded the datasets of the GTEx Project [31] to inspect the gene expression of these genes in different tissues, as shown in Fig. 4. Among the 12 genes selected by GenEpi, 11 have high expression level in the brain tissues. In addition, five genes, *CLEC16A, VSNL1*, *SYT6*, *CACNA1E* and *LINC00299*, have a similar expression pattern with *APOE*. These 12 genes are categorized as cross-gene epistasis, single-gene epistasis and single-SNP features based on the feature types selected by GenEpi. GenEpi detected only one cross-gene epistasis, which is *MICB * TOB2*. We found several evidence to demonstrate that this interaction might have true association with AD. MIC proteins are induced on the cell surface by stress and are ligands of a common activatory natural killer-cell receptor (NKG2D) [32]. They thus have roles in antimicrobial defense, tumor suppression and autoimmune-like diseases [33]. Quiroga et al. has found that the human MHC class I chain-related genes (MICA and MICB) are located within the HLA region that has stratified association with AD in the cohort of the Oxford Project to Investigate Memory and Ageing (OPTIMA) even though the association did not survive the correction for multiple testing [34]. TOB2 has been reported as a most up-regulated gene [35] by expression analysis on multiple AD expression datasets from the GEO database to identify significant genes associated with electrophysiological pathways and attempted determination of interconnected canonical molecular pathways. The interaction of MICB and TOB2 suggests a not-yet-explored biological process in AD pathogenesis. However, the significant associations of these genes in genome-wide association studies associated with immune system diseases and neurological diseases (see GWAS Catalog, 22561518, 29083406, 28540026), warrant future investigation.

**Fig. 4.**
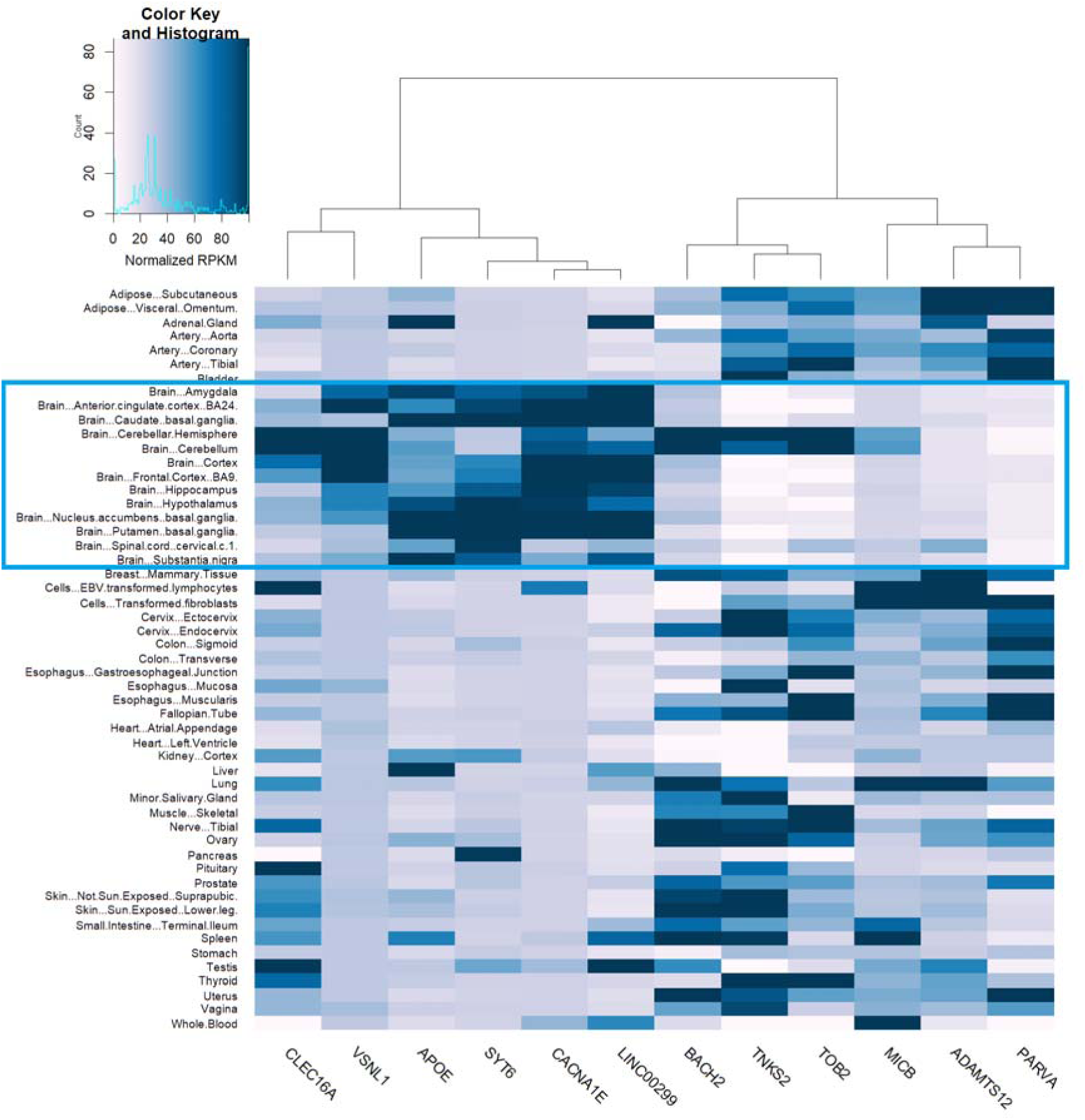
The heatmap of gene expression in different tissues for the 12 genes selected by GenEpi. The blue box highlights the sub-regions of brain.

About the 11 single-gene epistasis, there are several possible reasons accounting for intramolecular SNP-SNP interactions identified in this study. The first is a synergistic regulation of transcription, the second is a synergistic interaction between transcriptional and post-transcriptional regulation, and the third is an intramolecular SNP pair modulating the expression of two separate neighboring genes. Most of the single-gene epistasis selected by GenEpi can be explained by these three possible reasons and only two of the SNP-SNP interactions are not immediately clear at this moment. First: a synergistic regulation of transcription. For example, rs12095538 resides in the eQTL of STY6 and rs2774308 resides in a DNAse region of STY6. It is very possible that variants in both rs12095538 and rs2774308 synergistically affect the expression of STY6. SYT6 encodes protein control neurotransmitter release. It has been shown that the expression of SYT6 is significantly down regulated in AD [36], supporting the causative role of the inherited rs12095538 and rs2774308 haplotype in AD progression. Likewise, rs56233035 and rs3678 reside in the LD blocks containing the eQTL and DNAse region of CACNA1E, respectively. CACNA1E encodes a subunit of CaV1.2 calcium channel. Its expression level in reactive astrocytes is associated with amyloid-β plaques formation in an AD mouse model [37]. Another examples are the interactions between rs12926153 and rs12922908, and rs9652600 and rs12922908 that are within DNAse regions controlling the expression of CLEC16A in brain. CLEC16A encodes a lectin receptor involved in inflammation process and is differentially expressed in AD brains [38]. Lastly, rs2052573 and rs34580133 reside in DNAse regions within the LINC00299 and thus may synergistically control the expression of this long non-coding RNA that has been implicated in brain development [39].

Second: a synergestic interaction between transcriptional and post-transcriptional regulation. For example, rs9344977, in an intron of BACH2 containing cis-regulatory elements, may affect the expression of BACH2, whereas rs56148686 in another intron of BACH2 without any cis-regulatory element. We suspect that rs56148686 may interact with s9344977 through a post-transcriptional mechanism such as RNA stability, RNA splicing, or microRNA binding. BACH2 encodes a transcription factor involving cellular responses toward oxidative stress. Interestingly, BACH2 is upregulated in cultured human neuroblastoma cells upon exposure to Alzheimer Amyloid β [40]. Similar mechanisms may apply to interaction pairs rs12189429-rs6881360 and rs12187423-rs6881360 that presumably regulating the abundance of ADAMTS12 mRNA. The importance of ADAMTS12 in AD is supported by another independent study using same ANDI dataset [41]. Third: an intramolecular SNP pairs modulating the expression of two separate neighboring genes. For example, rs11675339 and rs2710687 of VSNL1 appear to control the expression of GEN1 and SMC6 respectively. We also found that VSNL (also known as VILIP-1) has strong evidences to AD [42–44]. GEN1 and SMC6 play pivotal roles in repairing double-strand breaks of genomic DNA. Interestingly, rDNA instability, a result of failures to repair double-strand breaks of genomic DNA, has been implicated in AD [45]. Most of the single-gene epistasis selected by GenEpi can be explained by these three possible modes and only two of the significance SNP-SNP interactions are not immediately clear at this moment. For example, rs12366151 resides in the eQTL of MICALCL, an AD associated gene [46] but the interacting rs10831829 resides in the intron of PARVA, a gene not-yet-reported to be associated with AD. Similarly, rs2421701 and rs200512701 reside in two eQTLs regulating TNKS2 expression. However, the ADP-ribose polymerase functionally contributes to AD is not clear.

Last, there are only two single-SNP features and both of them are located in APOE, which is a well-known causal gene of AD, revealing that GenEpi is an effective tool to identify disease-causing genes. Moreover, GenEpi successfully selected out the SNP rs429358, which determines the allele type of APOE with rs7412. While GenEpi has shown its ability to identify epistasis efficiently, it might still has the following limitations. Firstly, GenEpi can only detect pairwise interactions. Second, GenEpi is a memory-consuming package, which might cause memory errors when calculating the epistasis of a gene containing a large number of SNPs. We recommend that the memory for running GenEpi should be over 256 GB. Finally, a small sample size may lead overfitting, which forces us to use strict thresholds during feature selection. In this way, GenEpi delivers a high precision rate, but might suffer having false negatives. This implies different GWAS data might detect different sets of true positives. In summary, the results of this study demonstrated that GenEpi is a promising software package to identify causal SNPs and epistasis in GWAS, and it can be further used to predict the phenotypes. With the demonstrated efficiency, GenEpi is a powerful tool to explore gene-gene interactions that underlie complex diseases.

## 4. Materials

This study applied GenEpi on an AD cohort, which was used in Alzheimer’s disease Dream Challenge [28], In total, the cohort consists of 767 participants, who were healthy elderly, mild cognitive impairment (MCI) and AD patients from the Alzheimer’s Disease Neuroimaging Initiative (ADNI) database. The 767 ADNI participants consist of 241 cognitively normal (CN), 130 Early MCI (EMCI), 273 Late MCI (LMCI) and 123 AD participants. We adopted only genetic features in this study. All the genetic data has been pre-processed by the organizers that held the challenge [28]. The genetic data were genotyped using the Illumina Human610-Quad BeadChip and Illumina HumanOmniExpress BeadChip. The multidimensional scaling analysis was applied by PLINK using HAPMAP3 to ensure that samples are within the cluster of European populations. Subsequently, the data were imputed according to the 1,000 genome haplotypes. After imputation, there were 12,809,667 genotype features in total. For predicting the diagnosis of AD, we used 364 participants, of which the clinical diagnosis are CN or AD, to predict which samples are control subjects or the AD patients.

## 5. Methods

The architecture of GenEpi is shown in Fig. 5. GenEpi is designed to group SNPs by gene boundaries. GenEpi first considers the genetic variants in a gene as features, which have a higher chance to interact with each other and to influence molecular functions. The idea of within-gene epistasis analysis followed by cross-gene analysis is not new, which has also been used in previous studies [47–50]. Differently, GenEpi adopts two-element combinatorial encoding when producing features and models them by L1-regularized regression with stability selection, which will be explained in Section 2.2. In the first stage (STAGE 1) of GenEpi, the genotype features from each single gene will be combinatorically encoded and modeled independently by L1-regularized regression with stability selection. In this way, we can estimate the prediction performance of each gene and detect within-gene epistasis with a low false positive rate. In the second stage (STAGE 2), both of the individual SNP and the within-gene epistasis features selected by STAGE 1 are pooled together to generate cross-gene epistasis features, and modeled again by L1-regularized regression with stability selection as STAGE 1. Finally, the user can combine the selected genetic features with environmental factors such as clinical features to build the final prediction models. In addition to the main procedures, two pre-processing steps are also implemented in GenEpi: retrieving the gene information from public databases and reducing the gene information from public databases and reducing the dimensionality of the features using linkage disequilibrium (LD). In the end, we released a Python package that implements GenEpi. The details of these steps and the GenEpi method will be described in the following sections.

**Fig. 5.**
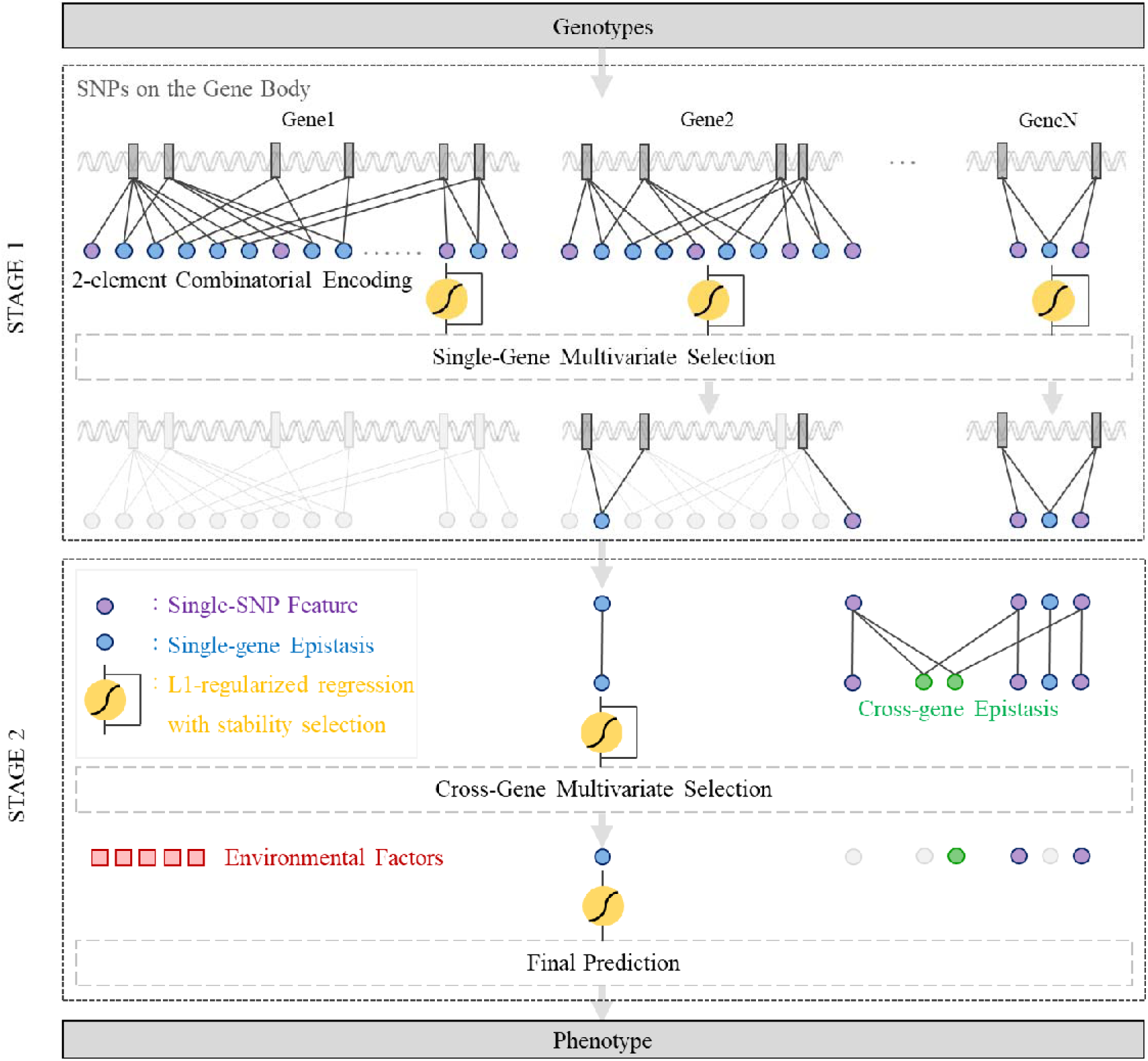
The architecture of GenEpi.

### University of California Santa Cruz (UCSC) database

To obtain the gene information such as official gene symbols and genomic coordinates, we retrieved kgXref and knownGene data table from the UCSC human genome annotation database [51, 52]. The version of the database we used is the Feb. 2009 assembly of the human genome hg19, GRCh37 Genome Reference Consortium Human Reference 37. The two data tables were merged in order to generate a local database containing the gene symbols as well as the genomic coordinates of each gene. The in-house script we built could update this local database automatically. It is noted that there are many different categories of genes in the RefSeq database. In this study, we only focused on the mRNA and non-coding RNA (22,376 genes in total). The selected transcripts were projected on the genomic coordinates and the coordinates of corresponding genes were determined based on the leftmost and rightmost positions of the corresponding transcripts. Moreover, to discover the factors that might affect the transcription of genes, we also retained the promoter region of each gene. In genetics, the promoter region is a segment of DNA that initiates the transcription of a particular gene. Promoters are located near the transcription start sites of genes, on the upstream of the same DNA strand (towards the 5’ region of the sense strand of the transcript). In general, a promoter region can be 100-1000 base pairs long. In this study, we extracted 1,000 nucleotides on the upstream of the starting position of each gene as the promoter region.

### Estimation of linkage disequilibrium

In GWAS datasets, a SNP often exhibits high dependency with its nearby SNPs because of linkage disequilibrium (LD). In the practical implantation, we prefer to group these dependent features to reduce the dimension of features. In other words, we can take the advantages of LD to reduce the dimensionality of SNP features. In this regard, we adopted the same approach developed by Lewontin [53] to estimate LD. We used D’ >0.9 and r^2^>0.9 as the criteria to group highly dependent SNP features as blocks. In each block, we chose the features with the largest minor allele frequency to represent other features in the same block.

### Discovery of within-gene epistasis

The main objective of the first stage in GenEpi is to select candidate features from each gene. In order to extract SNP features for a gene, we used the start and end positions of each gene from the local UCSC database to split the SNP features after dimension reduction. Since there are 22,376 genes in the UCSC database, we obtained 22,376 subsets of the SNP features. In each subset, A SNP feature with the alleles ‘A’ and ‘a’ could have three possible genotypes, AA, Aa and aa, which are used to refer to the pairs of alleles. The pairs of alleles are subsequently separated into three binary features using one-hot encoding. In order to evaluate epistasis, we generated interacting features by crossing each pair of genotype features. Considering the false positive rate and computational complexity, we only focused on pairwise interactions of epistasis throughout this study. We defined the interaction between two SNPs in Equation 1. In Equation 1, ***α*_1_*SNP*_1_** + ***α*_2_*SNP*_2_** stand for the additive interactions and ***α**_int_*_(1,2)_***SNP*_1_⊗*SNP*_2_** represents the synergistic interactions that contain nine terms.

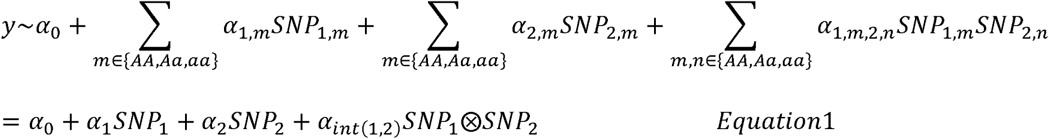

Before modeling each subset of genotype features, two criteria were adopted to exclude low quality data. The first criterion is that the genotype frequency of a feature should exceed 5%, where the genotype frequency means the proportion of genotype among the total samples in the dataset. The second criterion is regarding the association between the feature and the phenotype. We used χ^2^ test to estimate the association between the feature and the phenotype, and the p-value should be smaller than 0.01. In the end, a gene may have multiple SNPs. The general form of the linear model for a gene with *k* SNPs is defined as Equation 2, which is termed as two-element combinatorial encoding.

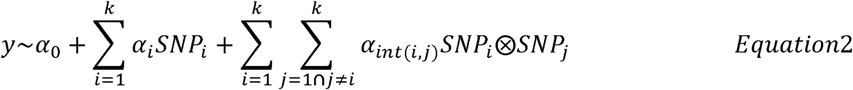

We conducted L1-regularized regression with stability selection [54] for modeling each gene. The first advantage is that the sparsity of the L1- regularized model prefers solutions with a smaller number of features, which effectively reduces the effect of feature dependency. The second advantage is by using stability selection [54], we can largely reduce the false positive rate. Stability selection works by resampling and remodeling the training set hundreds of times, followed by picking out the features that are repeatedly selected across randomization. In this study, we executed this randomization 500 times, and the features selected by stability selection would be retained for the next stage.

### Discovery of cross-gene epistasis

In the second stage, we used the features selected by STAGE 1 to generate cross-gene epistasis features and then applied the same selection procedure described in Section 2.3 to find the cross-gene epistasis that are associated with the phenotype. The procedures were slightly modified here. Since we only focused on pairwise interactions, instead of using the entire features we selected in STAGE 1, we only used single-SNP features to generate cross-gene epistasis features. Also, we used the genotype frequency and the p-values of χ^2^ test to control the quality of features and to avoid overfitting. Nevertheless, the p-value of each feature in this stage should be smaller than 10-5. All of the features from different genes would be merged for modeling cross-gene epistasis. We conducted L1-regularized regression for modeling, and the stability selection were used once again to select the final genotype feature set. Since the phenotype may also be affected by environmental factors, after determining the final set of genotype features, the user can included the environmental factors such as clinical assessments for constructing the final model. Subsequently, the final model was evaluated through a process called double cross validation (CV). In the external loop of double CV, all the instances were divided into two subsets to serve as training and independent test sets. In this study, we used 2-fold CV and leave-one-out CV (LOO CV) in external loop for evaluation. In the internal loop, we also used 2-fold CV for model selection.

## Declarations

### Ethics approval and consent to participate

Not applicable.

### Consent for publication

Not applicable.

### Availability of data and material

Not applicable.

### Authors’ contributions

Y.-C.C. initiated the study, designed the analysis procedures, performed the analysis and wrote the manuscript. J.-T.W. finished the literature survey of the epistasis that selected by GenEpi and wrote the discussion. C.-Y.C. is the main advisor of Y.-C.C., guided this research on right way and provided ideas to optimize the methods. Y.-C.C, M.-Y.H., Y.-A.T, C.-Y.C. and Y.-J.O. were team members of Alzheimer’s disease Dream Challenge. C.-Y.C., Y.-J.O., P.-H.H. K.M.G. and S.W.Y. commented on the draft and revised the manuscript. All authors read and approved the final manuscript.

## Acknowledgements

Data collection and sharing for this project was funded by the ADNI (National Institutes of Health Grant U01 AG024904). ADNI is funded by the National Institute on Aging, the National Institute of Biomedical Imaging and Bioengineering, and through generous contributions from the following: Abbott; Alzheimer’s Association; Alzheimer’s Drug Discovery Foundation; AmorfixLife Sciences Ltd.; AstraZeneca; Bayer HealthCare; BioClinica, Inc.; Biogen Idec Inc.; Bristol-Myers Squibb Company; Eisai Inc.; Elan Pharmaceuticals Inc.; Eli Lilly and Company; F.Hoffmann-La Roche Ltd and its affiliated company Genentech, Inc.; GE Healthcare; Innogenetics, N.V.; Janssen Alzheimer Immunotherapy Research & Development, LLC.; Johnson & Johnson Pharmaceutical Research & Development LLC.; Medpace, Inc.; Merck & Co., Inc.; Meso Scale Diagnostics, LLC.; Novartis Pharmaceuticals Corporation; Pfizer Inc.; Servier; Synarc Inc. and Takeda Pharmaceutical Company. The Canadian Institutes of Health Research is providing funds to support ADNI clinical sites in Canada. Private sector contributions are facilitated by the Foundation for the National Institutes of Health (www.fnih.org). The grantee organization is the Northern California Institute for Research and Education and the study is coordinated by the Alzheimer’s Disease Cooperative Study at the University of California, San Diego. ADNI data are disseminated by the Laboratory for Neuro Imaging at the University of California, Los Angeles. This research was also supported by NIH grants P30 AG010129, K01 AG030514 and the Dana Foundation.

